# Riverscape genetics in brook lamprey: genetic diversity is less influenced by river fragmentation than by gene flow with the anadromous ecotype

**DOI:** 10.1101/866533

**Authors:** Quentin Rougemont, Victoria Dolo, Adrien Oger, Anne-Laure Besnard, Dominique Huteau, Marie-Agnès Coutellec, Charles Perrier, Sophie Launey, Guillaume Evanno

## Abstract

Understanding the effect of human induced landscape fragmentation on gene flow and evolutionary potential of wild populations has become a major concern. Here, we investigated the effect of riverscape fragmentation on patterns of genetic diversity in the freshwater resident brook lamprey (*Lampetra planeri*) that has a low ability to pass obstacles to migration. We also tested the hypotheses of i) asymmetric gene flow following water current and ii) admixture with the closely related anadromous *L. fluviatilis* ecotype having a positive effect on *L. planeri* genetic diversity. We genotyped 2472 individuals, including 225 *L. fluviatilis*, sampled in 81 sites upstream and downstream from barriers to migration, in 29 West-European rivers. Linear modelling revealed a strong positive relationship between the distance to the source and genetic diversity, consistent with expected patterns of decreased gene flow into upstream populations. However, the presence of anthropogenic barriers had a moderate effect on spatial genetic structure. Accordingly, we found evidence for downstream-directed gene flow, supporting the hypothesis that barriers do not limit dispersal following water flow. Downstream *L. planeri* populations in sympatry with *L. fluviatilis* displayed consistently higher genetic diversity. We conclude that genetic drift and slight downstream gene flow mainly drive the genetic make up of upstream *L. planeri* populations whereas admixture between ecotypes maintains higher levels of genetic diversity in *L. planeri* populations sympatric with *L. fluviatilis*. We discuss the implications of these results for the design of conservation strategies of lamprey, and other freshwater organisms with several ecotypes, in fragmented dendritic river networks.

## Introduction

Human activities strongly modify natural ecosystems (Vitousek *et al.*, 1997) and impact evolutionary trajectories of wild species (Palumbi, 2001) posing unprecedented threats on the maintenance of biodiversity (Ceballos *et al.*, 2017). In particular, habitat fragmentation is a major threat to wild species (Vitousek *et al.*, 1997; Fahrig, 2003). Habitat fragmentation can reduce gene flow among subpopulations, which can ultimately decrease effective population size and genetic diversity (Frankham, 1998; Couvet, 2002; DiBattista, 2008; Blanchet *et al.*, 2010; Whiteley *et al.*, 2013). Habitat fragmentation influences the genetic structure and diversity of various species including birds (Alonso *et al.*, 2009), fishes (Hänfling and Weetman, 2006; Blanchet *et al.*, 2010; Torterotot *et al.*, 2014; Gouskov *et al.*, 2016) and plants (Young *et al.*, 1996).

Small isolated populations are expected to fix weakly deleterious alleles by random drift (Lynch *et al.*, 1995; Glémin, 2003) and can suffer higher extinction risks (Saccheri *et al.*, 1998; Carlson *et al.*, 2014; Smith *et al.*, 2014)). Maintaining high connectivity and genetic diversity levels is thus fundamental to preserve the evolutionary potential of populations (Frankham *et al.*, 2014; Frankham, 2015; Ralls *et al.*, 2018). Freshwater ecosystems have been particularly affected by fragmentation worldwide (Dynesius and Nilsson, 1994; Nilsson *et al.*, 2005) due to the construction of dams, weirs, and to artificial modifications of river channels. Such fragmentation alters the possibility of gene flow between populations of aquatic organisms, so that upstream isolated populations are particularly exposed to genetic drift and its consequences, namely reduced genetic diversity and ultimately increased inbreeding. In addition, river systems are naturally shaped as dendritic networks where migration preferentially occurs following downstream directed water flow, generating patterns of asymmetric gene flow (Hänfling and Weetman, 2006; Pollux *et al.*, 2009). As a result, populations are structured following a source-sink model (Kawecki and Holt, 2002; Hänfling and Weetman, 2006) in which the genetic diversity will be smaller in upstream source populations than in downstream sink populations. Three possible processes may explain this pattern of downstream increase in genetic diversity observed across taxa (Paz-Vinas *et al.*, 2015): *i)* downstream biased dispersal generating downstream gene flow (Paz-Vinas *et al.*, 2013), *ii)* increase in downstream habitat availability (e.g. (Raeymaekers *et al.*, 2008) and *iii)* upstream founding events with loss of genetic diversity e.g. following postglacial colonization (Cyr and Angers, 2011). However, it remains unclear how human mediated alterations of habitat connectivity in rivers may obscure or exacerbate this pattern.

To date, most studies focused on delineating the effect of barriers to migration in large species targeted by fisheries. This is particularly the case for Salmonids that display a strong migratory behaviour and a good ability to pass obstacles (Morita and Yamamoto, 2002; Yamamoto *et al.*, 2004). In contrast, few empirical studies have focused on species with low dispersal abilities or weak capacities to pass obstacles (e.g. (Raeymaekers *et al.*, 2008; Blanchet *et al.*, 2010), which are expected to be more impacted by the effect of river fragmentation. In addition, species can display various life history strategies, which may differ in their dispersal capacity and thus be differentially affected by changes in habitat connectivity. For instance, in certain fish species some individuals are freshwater-resident whereas others are anadromous (i.e. reproduce in freshwater and juveniles migrate to sea for growth) (Jonsson and Jonsson, 1993; Dodson *et al.*, 2013). Anadromous individuals can either display a homing behavior as they return back to their natal river to spawn, or disperse into neighboring rivers, which can enhance gene flow. Consequently, anadromous populations generally display lower levels of population genetic structure than resident populations (Hohenlohe *et al.*, 2010; Spice *et al.*, 2012; Hess *et al.*, 2013; Rougemont *et al.*, 2015, 2017). It has also been shown that anadromous salmonid populations usually display a higher level of genetic diversity than resident populations (e.g. (Perrier *et al.*, 2013)but it is not clear whether admixture between both forms may enhance genetic diversity of resident populations when both forms coexist.

The European brook lamprey *Lampetra planeri* is a widespread freshwater resident species with a putatively low dispersal ability linked to its small size (15-22 cm) and its particular life cycle, as the adults live only a few months in the river before they reproduce and subsequently die (Taverny and Élie 2010). It is closely related to the river lamprey *L. fluviatilis* that is parasitic and anadromous at the adult stage. The two taxa share many similarities (e.g. juveniles spend 3 to 6 years burrowed in the substrate of river beds) and they are best described as partially reproductively isolated ecotypes (Rougemont *et al.*, 2017). However, the distribution of *L. fluviatilis* is often restricted to lower parts of rivers due to a low upstream migratory ability in the presence of obstacles (Lucas *et al.*, 2009; Kemp *et al.*, 2011; Russon *et al.*, 2011; Foulds and Lucas, 2013). *L. fluviatilis* populations from nearby watersheds remain connected and display a low genetic differentiation in relation to dispersal abilities through the marine environment and an apparent moderate homing behavior (Rougemont *et al.*, 2015; Bracken *et al.*, 2015). In contrast, *L. planeri* has a highly reduced migratory behavior: it does not move outside its watershed and generally migrates on short upstream distances within the river for breeding purposes (Malmqvist, 1980). Thus, the most isolated brook lamprey populations located in the upper reaches of rivers can be strongly genetically differentiated from other populations either downstream or in other rivers (Mateus *et al.*, 2011; Rougemont *et al.*, 2015; Bracken *et al.*, 2015). These isolated populations often display a low genetic diversity at microsatellite loci (Rougemont *et al.*, 2015). Besides, the dispersal ability of *Lampetra* larvae that spend 5 to 7 years in freshwater is largely unknown but downstream dispersal may be important at this stage. In particular, flood events producing the remobilisation of fine sediments where larvae are burrowed may favor passive drift of larvae. Such downstream dispersal should further enhance the natural tendency to increase in genetic diversity downstream river networks, hence we hypothesize that in brook lamprey gene flow should clearly be asymmetric. In addition, brook lamprey populations living in sympatry with river lampreys have been found to display a higher level of genetic diversity than populations located in upstream reaches where river lampreys are absent (Rougemont et al. 2015, 2016). As a result, brook lamprey populations may generally benefit from gene flow with *L. fluviatilis* that may act as a ‘reservoir’ of genetic diversity. However, to disentangle the effects of gene flow between ecotypes and downstream biased dispersal, the genetic diversity of *L. planeri* populations should be compared between rivers where only *L. planeri* is present and watersheds where both species coexist, which has not be tested yet.

The main aims of this study were to understand: i) the role of river fragmentation on population genetic diversity and structure of *L. planeri* in various river systems from North western Europe, ii) the extent of asymmetry in gene flow among *L. planeri* populations from the same river, and iii) the possible role of *L. fluviatilis* in increasing genetic diversity in sympatric *L. planeri* populations *via* introgression. We performed extensive sampling of *L. planeri* upstream and downstream barriers to migrations in 29 rivers from three hydrogeologic regions: Brittany, Normandy and Upper Rhône, in France. Moreover, two watersheds were sampled more extensively to further investigate the combined effects of multiple barriers to migration on patterns of genetic diversity. To test the prediction that *L. planeri* populations found in sympatry with river lampreys may display greater levels of genetic diversity than populations where river lampreys are absent, we sampled sympatric and parapatric populations of *L. planeri* in Normandy and populations in Brittany where river lampreys are absent.

## Materials and Methods

### Sampling design

In 2013 and 2014 we sampled with electrofishing 2502 lamprey individuals in each of 81 sites spread over 29 rivers (**Figure 1**). We targeted *L. planeri* located upstream and downstream of a putative barrier to dispersal and, if possible, close to the barrier (less than 1 km upstream or downstream) to limit the effect of isolation by distance. In this study a site thus corresponds to a sampled point located either downstream or upstream a barrier to migration. We considered all kinds of barriers of moderate size (height between 0.50 m and 5 meters, described in Supplementary Table S1) that may limit the dispersal of lampreys. Although we initially planned to include the age of barriers in our model, such data were rarely available. In some cases, we were unable to capture lampreys immediately downstream or upstream dams and some sampled points were separated by more than one obstacle. In addition, 12 pairs of sites from eight rivers without barriers to migration were included in the dataset to investigate the effect of distance alone. 2274 *L. planeri* lampreys were collected in 73 sites in four distinct regions from France (Mediterranean area (Rhône), Normandy, Brittany and Upper Rhine), as well as 3 sites in United Kingdom and Ireland. In the 17 sites (n= 536 individuals) from Normandy *L. planeri* coexist in sympatry or parapatry (i.e. populations from the same river separated by obstacles to migration) with *L. fluviatilis*. Conversely, in the 32 sample sites from Brittany *L. planeri* (n= 969 individuals) are allopatric since no *L. fluviatilis* are currently present in these coastal rivers. In addition, two sites (one upstream and one downstream a migratory barrier) were sampled from a tributary of the Loire (the Cens River). In Brittany, for two rivers we failed to capture *L. planeri* both upstream and downstream a barrier, so only 30 sites were suitable to study the effect of fragmentation in this area. We also sampled 18 sites (n= 575 individuals) from the Rhône area and Upper Rhine to better capture the geographic distribution of genetic diversity. In addition, 228 *L. fluviatilis* were sampled in 8 populations: 7 from Normandy and one from the Loire River.

**Figure 1 :**
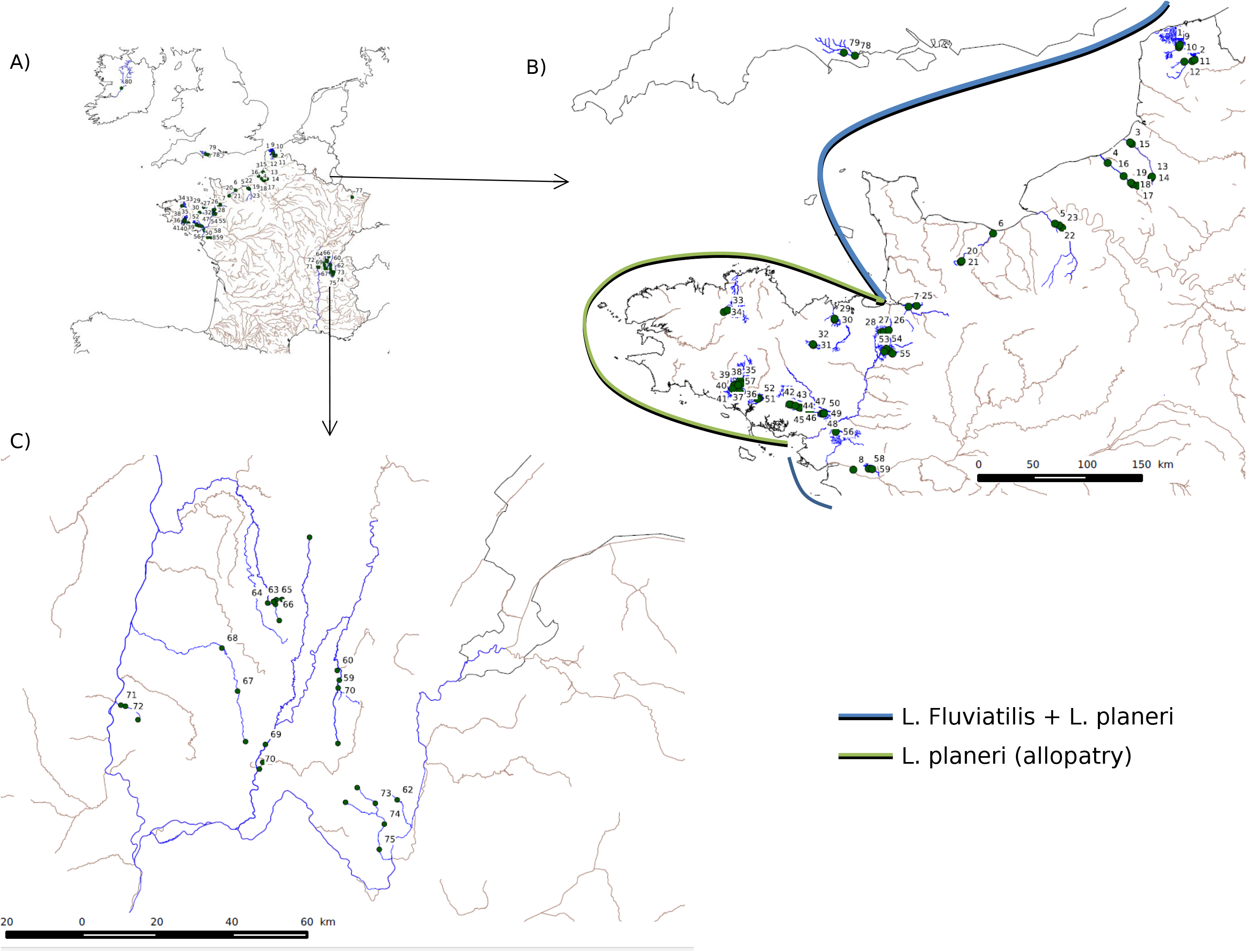
Sampling Map. A) Overview of the sampling site B) Zoom in the Normandy and Britanny diplsaying Allopatric, Parapatric and Sympatric Areas C) Zoom in the Upper Rhone, composed of Allopatric site only

A fin was clipped on each specimen and preserved in 95% EtOH. Explanatory variables of genetic parameters included the number of obstacles, their cumulated height, the geographic distances between each sample point and the distance to the river source. Data about obstacle height were gathered from the French “Referentiel des Obstacles à l’Ecoulement” (available at: http://carmen.carmencarto.fr/66/ka_roe_current_metropole.map). Geographic distances were computed using QGis 2.10.1. In addition, we performed linear transects in two independent rivers (the Arz and Scorff) including 8 and 7 sample sites respectively, to investigate the respective effects of obstacles and isolation by distance (Figure S2).

### Microsatellite genotyping

Genotyping was performed with 13 microsatellite markers specifically developed for *L. planeri* and *L. fluviatilis* after DNA extraction using a Chelex protocol modified from Estoup (1996) and strictly following protocols of (Gaigher *et al.*, 2013; Rougemont *et al.*, 2015)

### Genetic diversity within populations

We tested deviations from Hardy-Weinberg equilibrium using Genepop 4.1.0 (Rousset, 2008) exact tests with Bonferroni corrections (Rice, 1989) α = 0.05) and computed the inbreeding coefficient (*Fis*) for each population using Fstat 2.9.3 (Goudet, 1995). Genetic diversity indices were computed and included the number of allele (An), Allelic richness (*Ar*), observed heterozygosity (*Hobs*) and expected heterozygosity (*Hnb*) using Fstat 2.9.3 (Goudet, 1995) and Genetix 4.05.2 (Belkhir *et al.*, 2004). We tested for significant differences in levels of genetic diversity (Ar, Hnb, and relatedness) as a function of “geographicalconnectivity” using Generalized Lineard Models in R with gaussian familiy. We considered fives levels of connectivity: highly connected (corresponding to *L. fluviatilis)*, connected (*L. planeri* in sympatry *with L. fluviatilis*), moderately connected *(L. planeri* in parapatry), disconnected (*L. planeri* in allopatry in Brittany), strongly disconected (*L. planeri* in the Upper Rhône).

### Genetic structure among populations

We computed Weir & Cockerham’s estimator of *F*_ST_ (Weir and Cockerham, 1984) between all pairs of populations and used permutations tests with Bonferroni corrections to test for significance in Fstat. We tested for global pairwise differences in *F*_ST_ between upstream and downstream sites and among the three major regions using permutations tested in Fstat (10,000 to 15,000 test in each cases) as well as pairwise t.test adjusted for multiple testing using FDR corrections in R (R Core Team, 2015). However, populations are expected to deviate from migration drift equilibrium and to show a downstream increase in genetic diversity resulting in biased *F*_ST_ that may reflect this gradient effect rather than true differences. As a result, we also used indices of genetic differentiation that are independent from variations in genetic diversity among populations: the Jost D (Jost, 2008)nd Hedrick G’st (Hedrick, 2005). We illustrated the distribution of pairwise *F*_ST_ values in R using the heatmap.2 function implemented in the gplots package.

The Bayesian clustering program Structure 2.3.3 (Pritchard *et al.*, 2000) was used to evaluate the number of clusters (*k*) within rivers with multiple sampling sites (ie. Crano & Arz Rivers). A clustering analysis performed with the whole data set is provided as supplementary material. 10 independent replicates per *k* value were performed. Markov Chain Monte Carlo simulations (MCMC) used 200 000 burn-in and 200 000 iterations under the admixture model with correlated allele frequencies (Falush *et al.*, 2003). We used log likelihood Ln Pr(X|K) and the ΔK method (Evanno *et al.*, 2005) to determine the most likely number of clusters in Structure harvester (Earl and vonHoldt, 2012). Plots were drawn using Distruct 1.1 (Rosenberg, 2004).

### Isolation by distance

First, we tested for a pattern of isolation by distance (IBD), globally in each major geographic area separately. Given the complete disconnection between the Rhône drainage (in terms of waterway distance) and the rest of the sampled sites we did not compute IBD over the whole dataset. We computed Mantel tests using the linearised distance F_ST_/(1-F_ST_) against the waterway geographic distance between all sample sites using the R package Vegan, with the Mantel statistic being based on Spearman’s rank correlation rho (Oksanen *et al.*, 2015).

### Effect of river fragmentation on genetic diversity and differentiation

The Allelic richness (*Ar) differential* was used as an estimator of difference in genetic diversity between upstream and downstream sites within each river. Independent variables included 1) the cumulated height of obstacles and 2) geographic waterway distance to the source. We initially also included the number of obstacles as independent variable, but it was highly correlated with the cumulated height (r = 0.845) hence we only kept the cumulated height for analyses. All distances were computed manually in Qgis following water flow. Given that more than two upstream/downstream sites were sampled in a number of rivers, the river was fit as a random factor. Similarly, given the very different patterns observed in the different geographical areas (presence or absence of lampreys, reduced diversity in the Rhône), we fitted region as a random factor. Models were tested using AIC as implemented in lme4 (Bates *et al.*, 2015) and car (Fox & Weisberg) packages in the R software (R Core Team, 2015). The significance of each variable was computed using a type 3 anova. Pseudo R^2^ were then calculated using the function r.squaredGLMM implemented in the package MuMIn (Barton, 2019).

Next, we tested the effect of barriers to gene flow on genetic differentiation. We used the linearized genetic distance *F*_ST_/(1- *F*_ST_) (Rousset, 1997) between both sites in each river to test the effect of obstacles on genetic differentiation patterns. The exact same procedure as above with the exact same samples was performed implementing mixed linear model using distance and cumulated height as independent variables and river as well as region as random variables.

### Testing for downstream increase in genetic diversity

In addition, we tested the prediction of an increase of neutral genetic diversity downstream (Raeymaekers *et al.*, 2008; Blanchet *et al.*, 2010; Paz-Vinas *et al.*, 2013), which we expected to be strong due to downstream biased dispersal, and more important in sympatry areas due to gene flow between ecotypes. We used point estimates of *Ar* in linear mixed models and included all upstream and downstream sites from all rivers. The distance to the source was used as predictor variable (fixed effect) while the river and region were considered as random effects. The two other variables, namely number of obstacles and cumulated height to the source were all highly correlated with the distance to the source, (r = 0.89 and r = 0.85) respectively and between each other (r = 0.956) and therefore not included in a single model but only tested separately. We also computed the pseudo R2 using the r.squaredGLMM function.

### Testing for asymmetric gene flow

The intensity and symmetry of recent migration between upstream and downstream populations was measured using BayesAss 1.3 (Wilson and Rannala, 2003). We used a total of 10 millions iterations, discarding the first 1 million as burn-in and sampled the MCMC every 1,000 intervals. Following the authors’ recommendations, we computed a rough 95% credible interval using the mean ± 1.96 std. We then considered that there was information in the data by considering only cases where the credible intervals of downstream and upstream-directed gene flow did not overlap. Then we assessed the symmetry of migration by normalizing the point estimates using (m_1 ← 2_ – m_2 ← 1_)/max[m_1 ← 2_,m_2← 1_] so that this index varies between −1 and +1. Here m_1 ← 2_ represents the fraction of individuals in population 1 (downstream) that are migrants from population 2 (upstream) each generation and m_2 ← 1_ represents migration in the reverse direction. Therefore, positive values indicate higher downstream directed migration whereas negative values indicate stronger upstream migration.

### Effect on local isolation by distance

Finally we measured IBD and tested to which extent it was affected by the presence of obstacles on the Crano and Arz River (Brittany region) where more sites had been sampled (7 and 8 sites respectively). We used Mantel and partial Mantel tests in R. We constructed matrices of linearized F_ST_ (G’_ST_ and D), Ar and Hnb differentials and matrices of pairwise waterway distances, number of obstacles and cumulated height for each river. Next, commonality analysis (Nimon *et al.*, 2008) was applied in order to take into account collinearity between distance and cumulated barrier heights between sampling sites. We assessed the extent to which each predictor variable contributed to the variance in the response variable *via* a set of unique and shared effects (Prunier *et al.*, 2015). The MBESS R package was used for this analysis (Kelley and Lai, 2012). at they did not provided additional information as compared to our linear models.

## Results

### Genetic diversity within populations

*Fis* was statistically significant (**Table S2**) in four populations: the downstream site on the Léguer River in Brittany (*F_is_* = 0.259) and three downstream sites in the Rhône watershed (Oignin *F_is_* = 0.598, Calonne *F_is_* = 0.495, Neyrieux *F_is_* = 0.395).

Levels of Allelic richness (*A_r_*) based on a minimal sample size of 11 varied from 1.18 (Reyssouze River, Upper Rhône) to 3.79 (Béthune River, Normandy) and from 1.20 to 3.85 for the mean number of alleles per locus (**Table 1, Table S2, Table S3**). Levels of *H_nb_* averaged over all loci per population also varied substantially, ranging from 0.011 (Reyssouze River) to 0.563 (Aa River, Normandy). Populations of the Upper Rhine displayed similar levels of diversity to those of Brittany (Table 1). On average *L. fluviatilis* populations were significantly more diverse than *L. planeri* populations both in terms of allelic richness (Table 1, p < 0.0042, 15 000 permutations) and expected Heterozygosity (**Table 1**, p < 0.0057, 15 000 permutations) (See also **Figure 2**). *L. Fluviatilis* populations in Normandy were not different from downstream *L. planeri* populations in Normandy in terms of expected heterozygosity or allelic richness (GLM, p > 0.05) (see **Table S4** and **Figure 2**). In contrast, the genetic diversity of *L. fluviatilis* populations was systematically higher than the one of upstream *L. planeri* populations of Normandy (i.e. parapatric) and from the neighboring *L. planeri* populations of Brittany (all p < 0.05, see table S4 and **Figure 2**). Levels of genetic diversity of the Frome (UK) and Shannon (Ireland) populations (Table 1) where similar to those observed in Normandy.Comparisons among geographical areas revealed a significantly lower (GLM, p > 0.05) genetic diversity of *L. planeri* populations from the upper Rhône compared to Brittany and Normandy (**Figure 2, Table S4 for detailed p-values**).

**Figure 2 :**
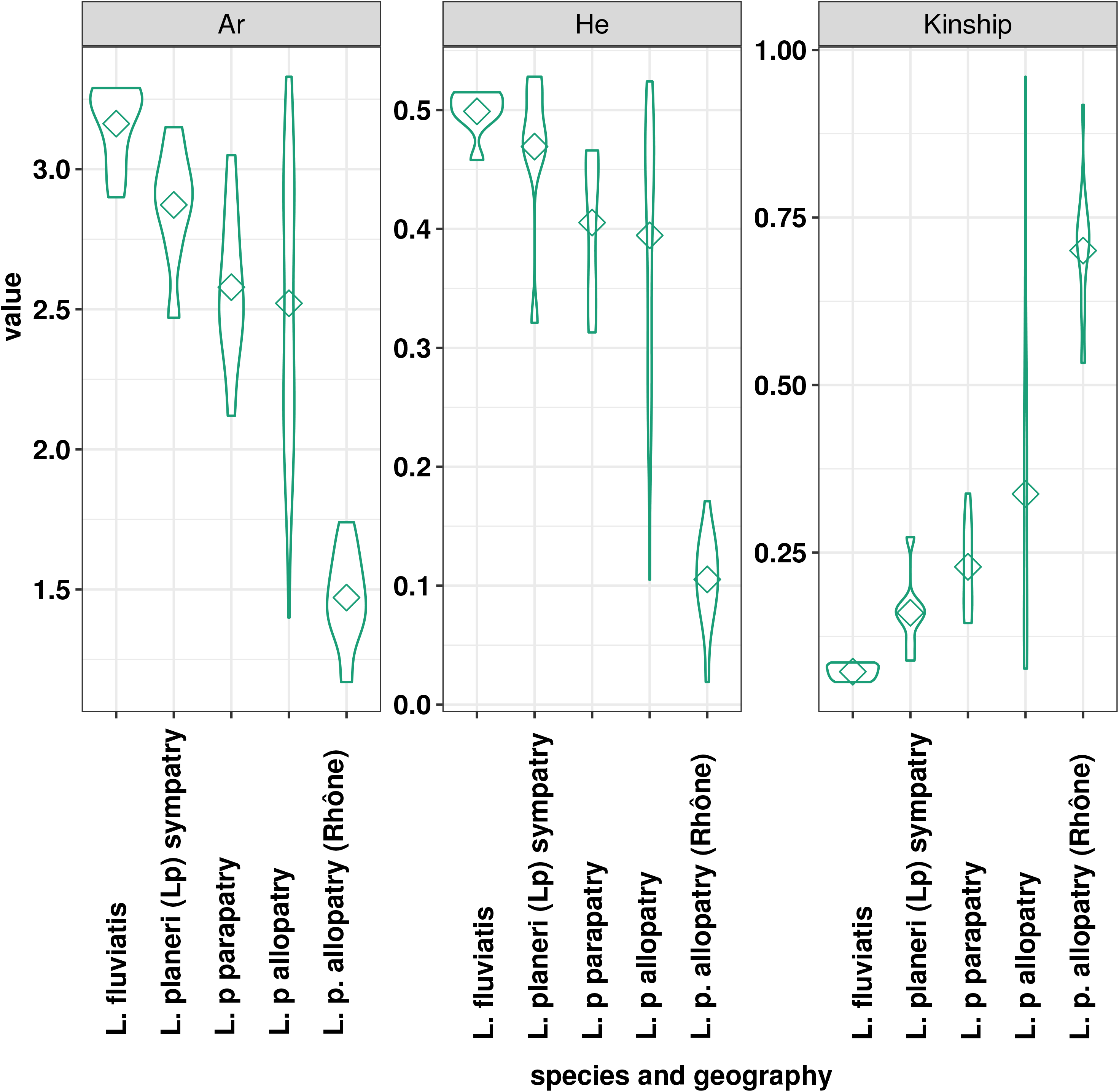
Violin plot of the distribution of genetic diversity and relatedness as a function of the geographic context and levels of connectivity. Diamond display median values. See text for statistical significance.

**Table 1:** Summary statistics of genetic diversity of *L. planeri* and *L. fluviatilis* populations for each geographic area. *N* = Number of individuals, *NbA* = Number of Alleles (averaged of all loci), Ar = Allelic Richness, *H*_e_ = unbiased Expected Heterozygosity, *H*_o_ = Observed Heterozygosity. *F_ST_* = Weir & Cockerham differentiation index and confidence interval computed in Fstat (Goudet, 1995).

|  | **N** | **NbA** | **Ar** | **He** | **Ho** | **$F_{ST}$ [95%IC]** |
| --- | --- | --- | --- | --- | --- | --- |
| Global | 2472 | 2.73 | 2.43 | 0.354 | 0.344 | 0.377 [0.334 - 0.418] |
| *L. fluviatilis*<br>(Normandy) | 225 | 3.84 | 3.39 | 0.505 | 0.491 | 0.003 [0 - 0.007] |
| *L. planeri* | 2247 | 2.65 | 2.37 | 0.344 | 0.334 | 0.396 [0.353 - 0.437] |
| Normandy. *L. planeri* | 536 | 3.15 | 2.88 | 0.435 | 0.440 | 0.139 [0.113 - 0.170] |
| Brittany& Normandy | 1505 | 2.94 | 2.62 | 0.416 | 0.411 | 0.241 [0.207 - 0.284] |
| Brittany | 969 | 2.88 | 2.58 | 0.406 | 0.396 | 0.279 [0.235 - 0.334] |
| Upper Rhône | 575 | 1.76 | 1.52 | 0.111 | 0.089 | 0.249 [0.047 - 0.317] |
| UK | 83 | 3.5 | 2.96 | 0.476 | 0.463 | NA |
| Ireland | 48 | 3.31 | 2.79 | 0.458 | 0.453 | NA |
| Upper Rhine | 36 | 3.00 | 2.58 | 0.355 | 0.323 | NA |

### Genetic differentiation and structure among populations

Global *F*_ST_ was 0.377 (95%IC = 0.334-0.418) and reached 0.394 (95%IC = 0.353-0.437) when excluding *L. fluviatilis* (**Table 1**). **The figure S1** illustrates two main groups of populations: the Upper Rhône *vs* all other populations. *L. fluviatilis* populations were weakly differentiated (*F*_ST_ = 0.005). Populations of *L. planeri* were significantly more differentiated than *L. fluviatilis* populations (p < 0.00017, 6000 permutations with Hierfstat after pooling). The highest *F*_ST_, equal to 0.90, was observed between the Reyssouze (Rhône) and the Moulin du Rocher sites (Brittany). The lowest *F*_ST_ was 0 as observed in several cases (**Figure S1** and **Table S5**). Average pairwise *F*_ST_ between upstream and downstream sites within a river was 0.025 [min = 0 – max = 0.095]. The maximal value of 0.095 was observed in the Crano between two sites located near the river source and in the absence of obstacles (**Table S5**). Pairwise *F*_ST_ between upstream and downstream sites were significant in 8 out of 43 pairwise comparisons with upstream-downstream sites from Normandy, where *L. fluviatilis* is present, being never significantly differentiated. Populations of the Upper Rhine, Frome and Shannon, were moderately differentiated from *L. fluviatilis* (**Table S5**). The Frome Downstream site in particular displayed modest differentiation from *L. fluviatilis* as it was not significantly different from the Hem, Risle and Oir river (*F*_ST_ below 0.0125). Finally, genetic differentiation between *L. planeri* populations in sympatry with *L. fluviatilis* (*i.e.* in Normandy) was lower than the averaged *F*_ST_ between *L. planeri* population living in allopatry from *L. fluviatilis* (Brittany). Results from analyses using both Hedrick *G*_ST_ and Jost D (data not shown) were largely similar to those based on Weir and Cockerham *F*_ST_.

### Population structure within rivers

Populations’ structure in the Crano (n=7 sites) and Arz River (n=8 sites) revealed similar patterns of admixture in these two systems. In the Crano, the two distinct uppermost tributaries formed distinct clusters with a lower degree of admixture than the downstream populations that displayed increased admixture values (**Figure S2**). In the Arz, the source population formed a slightly distinct cluster with lower admixture than the downstream populations (**Figure S2**). Details of population structure over the whole dataset are provided as supplementary materials (**Figure S3 and S4, Table S6)**.

### Landscape genetics

### Global Isolation by distance

First, we tested for global isolation by distance in each major region (Brittany, Rhône and Normandy). Mantel tests indicated a significant relationship between distance and linearized genetic differentiation in the upper Rhône area (Mantel r = 0.469, *p* = 2e^−4^). The pattern of IBD was less pronounced in Brittany, but still significant (Mantel r = 0.188, *p* = 0.016). In contrast, the pattern of isolation by distance was not significant in the Normandy area (Mantel r = 0.145, *p* = 0.143). This absence of relationship was largely driven by the lack of genetic differentiation between the upstream/downstream populations of the Oir River as compared to the remaining *L. planeri* Normandy populations. When this population was removed, the pattern of IBD appeared the strongest among Normandy populations (Mantel r = 0.55, *p* = 1e^−4^). To gain further insights about the evolutionary relationships among populations from coastal areas either connected to *L. fluviatilis* (Normandy) or disconnected (Brittany), we tested the pattern of IBD by keeping one (downstream) site per river. In this case the signal of IBD remained significant in Normandy (Mantel r = 0.43, *p* = 0.042) but not in Brittany (Mantel r = −0.0208, *p* = 0.53).

### Effect of river fragmentation on genetic diversity & differentiation

The model selection procedure based on AIC indicated that the best model included only the pairwise geographic distance (**Table 2**) with a highly significant effect on AR differential between downstream and upstream sites. Even though the cumulated height had a significant effect (*p = 0.002*) this model had the highest AIC (**Table 2**). The amount of variance of the best model explained by fixed factors was R2m = 0.21 whereas the entire model explained a greater part of the variance (pseudo R2c = 0.59). Regarding genetic differentiation, the Linear modelling approach revealed no significant effect of the tested variables (**Table 2**).

**Table 2:** Effect of landscape fragmentation on genetic diversity (AR differential) and genetic differentiation (*F*_ST_) between pairs of sites. Mixed linear models were used with river and region fitted as random factors. Model 1 = Distance + Barrier Size, Model 2 = Barrier Size Model 3 = Distance. AIC are provided for each model along with p-valeus and slope of the tested variables. R^2^m corresponds to the marginal R^2^ and represents the variance explained by fixed effects. R^2^c corresponds to the conditioned R^2^ and represents the variance explained by both fixed and random effects. Genetic differentiation was linearized using : Y = FST/(1- FST).

| Model | Effect of barrier on allelic richness |  |  |  |  |  | Effect on genetic differentiation |  |  |  |  |  |
| --- | --- | --- | --- | --- | --- | --- | --- | --- | --- | --- | --- | --- |
| | AIC | F | p | Slope | $R^2_m$ | $R^2_c$ | AIC | F | p | Slope | $R^2_m$ | $R^2_c$ |
| Model 1 : | -27.0 |  |  |  | 0.204 | 0.604 | -230.8 |  |  |  | 0.043 | 0.183 |
| Distance |  | 1.30 | **0.0003** | 0.008 |  |  |  | 0.993 | 0.322 | 0.0026 |  |  |
| Barrier size |  | 14.12 | 0.25 | 0.010 |  |  |  | 0.771 | 0.383 | 0.0005 |  |  |
| Model 2 : | -25.7 |  |  |  | 0.091 | 0.505 | -245.0 |  |  |  | 0.032 | 0.153 |
| Barrier size |  | 10.38 | **0.0018** | 0.026 |  |  |  | 2.615 | 0.110 | 0.0036 |  |  |
| Model 3 : | -35.3 |  |  |  | 0.212 | 0.586 | -241.9 |  |  |  | 0.036 | 0.188 |
| Distance |  | 26.45 | **0.000002** | 0.009 |  |  |  | 2.432 | 0.123 | 0.0008 |  |  |

### Effect of the distance to the source

As expected, we found a highly significant positive relationship between the distance to the source and the levels of genetic diversity (*p* = *0.000001***, F = 29.7)**. Testing each dependant variable revealed a significant effect of the barrier count and of the cumulated height (**Table 3**).

**Table 3:** Results of Mantel tests and Partial Mantel tests performed on the Arz and Crano rivers. Factors in brackets correspond to controlled effects in partial mantel tests.

|  | Arz |  | Crano |  | Arz |  | Crano |  |
| --- | --- | --- | --- | --- | --- | --- | --- | --- |
| | Y = $F_{ST} / (1 - F_{ST})$ | | | | Y = Allelic Richness Dif erential | | | |
|  | *r* | *p* | *r* | *p* | *r* | *p* | *r* | *p* |
| Dist | 0.163 | 0.237 | 0.86 | <0.001 | 0.72 | 6.00E-04 | 0.645 | 0.014 |
| N. obst | 0.018 | 0.273 | 0.31 | 0.152 | 0.62 | 0.002 | 0.33 | 0.133 |
| Height | 0.174 | 0.171 | 0.222 | 0.302 | 0.58 | 0.004 | 0.59 | 0.057 |
| Dist (N.obst) | 0.338 | 0.0357 | 0.885 | 0.0055 | 0.45 | 0.016 | 0.513 | 0.06 |
| Dist(Height) | 0.02771 | 0.389 | 0.799 | 0.024 | 0.52 | 0.006 | 0.379 | 0.155 |
| N.obst (Dist) | -0.300 | 0.901 | -0.77 | 0.965 | -0.067 | 0.623 | -0.0487 | 0.581 |
| Height (Dist) | 0.066 | 0.322 | -0.592 | 0.918 | -0.102 | 0.6487 | 0.222 | 0.266 |

### Downstream directed gene flow

Analysis of recent migration rates in BayesAss indicated that there was enough information in the data in 46% of the upstream-downstream comparisons (i.e. 18 out of 39 in which CI did not overlap, (**Figure 3, Table S7**). In 100% of these informative cases, we found that migration was predominantly directed from the upstream to the downstream areas with the index of asymmetry reaching a median value of 0.90 (average = 0.88). Furthermore, there was no difference in the intensity of gene flow between *L. planeri* pairs located in sympatry areas (Normandy) and *L. planeri* pairs in allopatry (Brittany and upper Rhône) (wilcoxon test W = 50, p = 0.117; GLM p> 0.1, **Table S8**), indicating that the highest genetic diversity observed in Normandy (**Figure 2**) was not due to a higher downstream directed gene flow in this area.

**Figure 3 :**
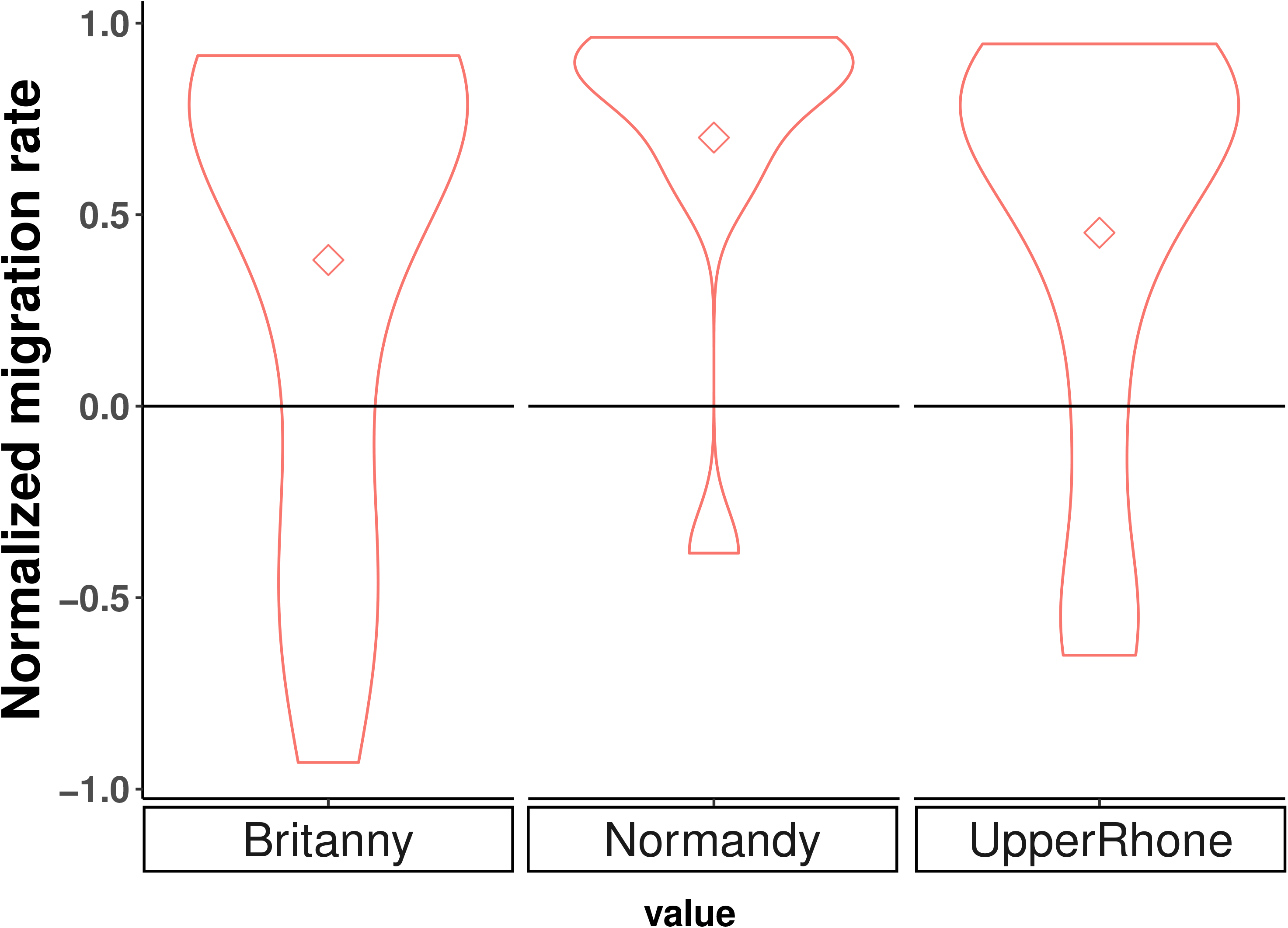
Violin plot of normalized estimates of recent migration rate (obtained with BayesAss) accross different geographic area displaying different levels of connectivity. Negative values indicate upstream directed migration. Positive values indicate downstream directed migration. The value of zero indicate symmetric migration. Diamonds correspond to the median value

### Effect on local isolation by distance

Mantel tests and partial Mantel tests on the two rivers where more than two sites had been sampled (Arz and Crano) indicated different influences of distance and obstacle-related variables. In the Arz River, all variables significantly influenced allelic richness (**Table 3)** whereas it was influenced solely by geographic distance in the Crano River. The extent of pairwise differentiation was also influenced by distance in the Crano River whereas this pattern was only revealed in the Arz when the influence of the number of obstacles was controlled for (**Table 3**). The commonality analysis **(Table 4)** also indicated a significant influence of the number of obstacles and geographic distance on genetic diversity (measured by heterozygosity) on the Arz with both contributions of unique and common effects, whereas only the number of obstacles influenced the pairwise differentiation. On the Crano River, commonality analysis indicated a strong influence (p < 0.01, **Table 4**) of the number of obstacles on pairwise differentiation whereas most of the variance in expected heterozygosity was explained uniquely by distance (**Table 4**).

**Table 4:**
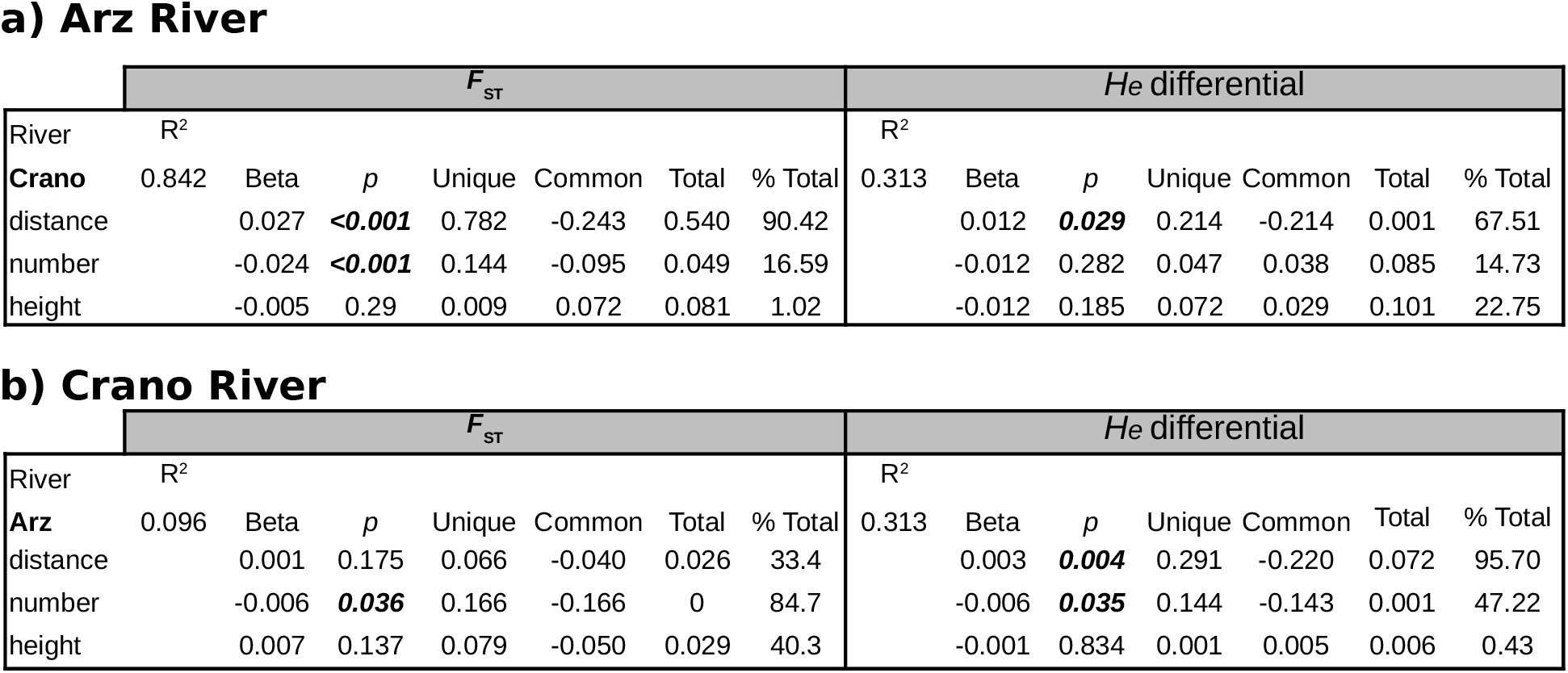
Commonality analyses performed on (a) the Arz River and (b) the Crano River. Unique = predictor unique effect Common = the sum of effects shared with other predictors Total = sum of unique and common contributions to the variance in the response variable.

## Discussion

The goals of this study were threefold: testing the effect of anthropogenic river fragmentation on patterns of population genetic diversity, testing for asymmetric gene flow and exploring the potential influence of the presence of *L. fluviatilis* on genetic diversity levels in *L. planeri.* We used *L. planeri* as a model to test the effect of fragmentation as this species displays a reduced migratory behaviour (Malmqvist, 1983). We found limited evidence for the effect of anthropogenic fragmentation on genetic diversity and differentiation of populations and the distance to the source was a more pertinent variable to explain patterns of genetic diversity within populations. Importantly, this lack of effect of river fragmentation could be explained by the strong downstream dispersal of *L. planeri*, which does not seem limited by obstacles of limited size considered in this study. The comparison of sympatric, parapatric and allopatric populations, located in downstream and isolated areas of different watersheds revealed a key role of *L. fluviatilis* in maintaining genetic diversity of *L. planeri* populations in the lower part of rivers where they co-occur.

### Small impact of anthropogenic fragmentation on the distribution of genetic diversity

While several studies have reported strong impacts of barriers to migration on either genetic diversity and/or structure (Hänfling and Weetman, 2006; Leclerc *et al.*, 2008; Raeymaekers *et al.*, 2008; Faulks *et al.*, 2010; Blanchet *et al.*, 2010; Torterotot *et al.*, 2014; Gouskov *et al.*, 2016), here evidence for such effects was low and factors such as the distance to the source or the distance between sites, strong downstream directed gene flow, all contributed to erase genetic differentiation and homogenize diversity levels. Our results are therefore slightly different from those of Bracken *et al.*, (2015) who suggested that barriers increased population differentiation. However, it seems that Bracken *et al.*, (2015) analysed separately the effects of distance and barriers, making the comparison difficult with our results.

The population genetic diversity was mostly affected by distance to the source, as upstream populations showed lower levels of allelic richness and heterozygosity (figure 4 and Table 5). This so called downstream increase in genetic diversity is expected in riverine habitat (Morrissey and de Kerckhove, 2009; Paz-Vinas *et al.*, 2015) and is frequently observed in empirical studies e.g. (Hänfling and Weetman 2006; Torterotot *et al.*, 2014; Gouskov *et al.*, 2015). Detailed investigations on the Arz River provided strong evidence for an increased downstream allelic richness and this pattern was also significantly influenced by all other physical variables. On the Crano, increase in genetic diversity was not influenced by geographic variables other than distance. A recent simulation study investigated the underlying processes that can generate this pattern (Paz-Vinas *et al.*, 2015), namely i) downstream-biased dispersal, ii) increase in habitat availability downstream and iii) upstream directed colonization. Among the three proposed processes, it appears likely that downstream dispersal plays a key role in *L. planeri* according to our analysis with BayesAss, which indicates higher upstream-downstream dispersal than downstream to upstream dispersal. Such dispersal is expected to occur passively given the long larval stage of lampreys. Indeed, larvae remain buried in the soft substrate of river beds for up to five years (Hardisty and Potter, 1971) and may passively be transported downstream (Dawson, *et al.*, 2015) during flood events. Bayesian clustering analysis (Fig S2) in the St-Sauveur-Crano river system (the Crano is a small stream flowing into the St Sauveur) revealed another important pattern explaining the increase in downstream genetic diversity *via* admixture among individuals originating from different upstream sites. The two upstream populations of the St Sauveur and Crano form two genetically distinct clusters (*F*_ST_ = 0.265) and individuals located downstream of the Crano appear admixed, possibly having a shared ancestry stemming from these two source populations (and possibly from other unsampled populations). The second process that may have generated low upstream genetic diversity is the occurrence of bottlenecks through multiple serial founder effects, following usptream river colonization after glacial retreats (Hewitt, 1996; Taberlet *et al.*, 1998). It remains unclear so far whether *L. planeri* populations have recovered from ancestral bottlenecks and disentangling the three hypotheses will require further data.

Overall, strong gene flow between upstream and downstream population likely explains our inability to detect the effect of river fragmentation globally. Moreover, even a small amount of upstream directed migration, as inferred here by BayeAss may contribute to reduce the effect of obstacles to migration. Alternatively, it is possible that subtle effects will be revealed later in time if most of the studied barriers are still relatively recent (Landguth *et al.*, 2010). Finer investigations on the Crano and the Arz revealed significant effects of distance and of the number of obstacles (according to the commonality analysis) on differentiation in the Crano River. In contrast, on the Arz River the effect of distance was only revealed when obstacles number was controlled for, in agreement with the commonality analysis. Finally, the impact of river fragmentation may be best revealed by study focusing on a single catchment and with bigger obstacles to migration (e.g. Raeymaekers *et al.*, 2008; Blanchet *et al.*, 2010; Gouskov *et al.*, 2015). We investigated the impact of obstacles of small to moderate size and it is possible that these obstacles do not influence the downstream passive drift of lamprey larvae, which may be sufficient to homogenize populations and obscure patterns of differentiation (Faubet *et al.*, 2007).

### River lamprey as a source of genetic diversity for resident lampreys

Understanding the evolutionary relationships between parasitic and nonparasitic lamprey ecotypes is a long-standing debate (Docker, 2009). Recent studies (Bracken *et al.*, 2015; Rougemont *et al.*, 2015) have shown that gene flow is ongoing between the river lamprey and the brook lamprey, locally lowering their level of genetic differentiation. Notably, Rougemont *et al.*, (2016, 2017) suggested the occurrence of locally asymmetric introgression from anadromous to resident sympatric populations following secondary contacts. Such introgression from a large marine population toward freshwater populations is also known to occur in the stickleback *Gasterosteus aculeatus (**Hohenlohe et al.*, 2010, 2012). Here, genetic analyses of populations in sympatry (on the same nest), in parapatry (where the two species cooccur on the same watershed but are geographically separated by impassable dams) and in allopatry (in coastal rivers where *L. fluviatilis* is absent) revealed that allopatric *L. planeri* displayed a lower genetic diversity than sympatric and parapatric populations (Figure 4). These results, with those from previous studies, further suggest that the current genetic makeup of *L. planeri* populations in Normandy is influenced by ongoing gene flow with *L. fluviatilis*. In addition, we found a much stronger pattern of IBD in the connected pairs of *L. planeri* (i.e. populations of downstream areas in Normandy) than in populations from Brittany. In the absence of inter-basin gene flow mediated by *L. fluviatilis*, populations of Brittany evolved independently from each other and do not seem globally at migration drift equilibrium (which does not imply that sub-populations within rivers deviate from this equilibrium). Populations from the Upper Rhône area displayed a highly reduced genetic diversity and were strongly differentiated from all other populations, which could be explained by different complementary hypotheses. First, there is evidence for at least three major evolutionary lineages existing in *L. planeri* (Espanhol *et al.*, 2007). It is thus possible that colonization of the Mediterranean area (Upper Rhône region) following postglacial colonization (the usual pattern in European fish species, (Bernatchez and Wilson, 1998)originated from a different lineage that the one having colonized the Atlantic and Channel areas. In these conditions, it is possible that our microsatellite marker set (originally developed using *L. planeri* and *L. fluviatilis* samples from the Atlantic and channel area) is not the most appropriate to perform accurate population genetic inference of Rhône samples. Second, *L. fluviatilis* no longer colonises this area and were already reported to be declining during the last century (Bernard, 1909; Gensoul, 1907). Consequently, it is possible that the history of divergence between Mediterranean and Atlantic populations was initiated a long time ago and that gene flow between neighbouring rivers of the Mediterranean area has been further reduced during the last century.

### Conservation implications

Fragmentation of rivers may impact lamprey populations, especially the most upstream populations that do not receive migrants from downstream sites. Whether the most isolated populations from headwaters suffer a mutation load and greater extinctions risks would require further investigations. It is now known from theory and empirical data that small isolated populations suffer greater inbreeding and extinction risks (Lynch, 1991; Higgins and Lynch, 2001; Spielman *et al.*, 2004; Frankham, 2005, 2015). On the other hand it is not clear if maintaining a possibility for upstream migration by removing obstacles may help preventing the loss of genetic diversity in the most uptream populations of *L. planeri* through the beneficial effects of gene flow (Frankham 2015). The source populations on the Crano and St Sauveur, as well as the high genetic differentiation on the Tamoute River despite the absence of migratory barrier clearly illustrate this issue.

Importantly, our study revealed positive impacts of the presence of *L. fluviatilis* on the maintenance of genetic diversity in sympatric populations. However, in Europe, *L. fluviatilis* abundance has strongly declined in some areas, due to habitat alteration and pollutions (Maitland *et al.*, 2015) and it is now considered as vulnerable in France in the IUCN red list (UICN France et al. 2019). In addition, the low ability of the anadromous ecotype to pass migration barriers often restricts its distribution to downstream areas where *L. planeri* are less abundant. In terms of conservation priority it appears fundamental to first ensure that *L. fluviatilis* will have access to upstream reaches of rivers. This will benefit both the river and brook lampreys in sympatric and parapatric areas. In such areas a joint regional management of the two ecotypes could be envisioned, whereas in allopatric areas, a management at the river scale may be more parsimonious.

### Conclusion

We have shown here that impacts of anthropogenic barriers to migration were modest on the extent of genetic differentiation, but we provided evidence that headwater populations of *L. planeri* displayed reduced genetic diversity, and were the most genetically differentiated as a result of isolation by distance in a river network. Despite the strong asymmetric gene flow toward downstream populations (probably due to passive drift of larvae), restoring the possibility for upstream active migration from downstream populations could increase genetic diversity and evolutionary potential in the most upstream populations (Brauer *et al.*, 2016; Pavlova *et al.*, 2017; Coleman *et al.*, 2018). In addition, our comparative analyses among sympatric, parapatric and allopatric areas support the hypothesis that sympatric populations display higher levels of genetic diversity due to introgression from *L. fluviatilis* (Rougemont *et al.*, 2015, 2016, 2017). Potential strong gene flow or even genome swamping from anadromous populations to resident populations thus plays a fundamental role in maintaining genetic diversity of *L. planeri* (Rougemont *et al.*, 2017). In addition, such introgression may play a key role through adaptive introgression and through the movement of freshwater adaptive alleles between adjacent rivers.

## Acknowledgements

We thank B. Jacquot, B. Herodet, B. Bulle, B. Dufour, Y. Salaville, H. Catroux, G. Sanson V. Zunigas, R. Pélerin, V. Mouren, M. Lesimple, B. Rigault, G. Garot, P. Domalain, J-L. Fagard, J.M. le Boucher, N. Jeanneau, A. Robbe, J. Tremblay (Experimental Unit of Aquatic Ecology and Ecotoxicology, U3E), Y. Perraud, P. McGinnity and B. Beaumont who helped us collecting samples. This study was funded by the Agence Française de la Biodiversité.

## Data Archiving

Raw genotype data used in this study will be available at the Dryad Digital Repository.

